## Supplementary material for "Evolution of imprinting via lineage-specific insertion of retroviral promoters": Bogutz-Supplementary Material

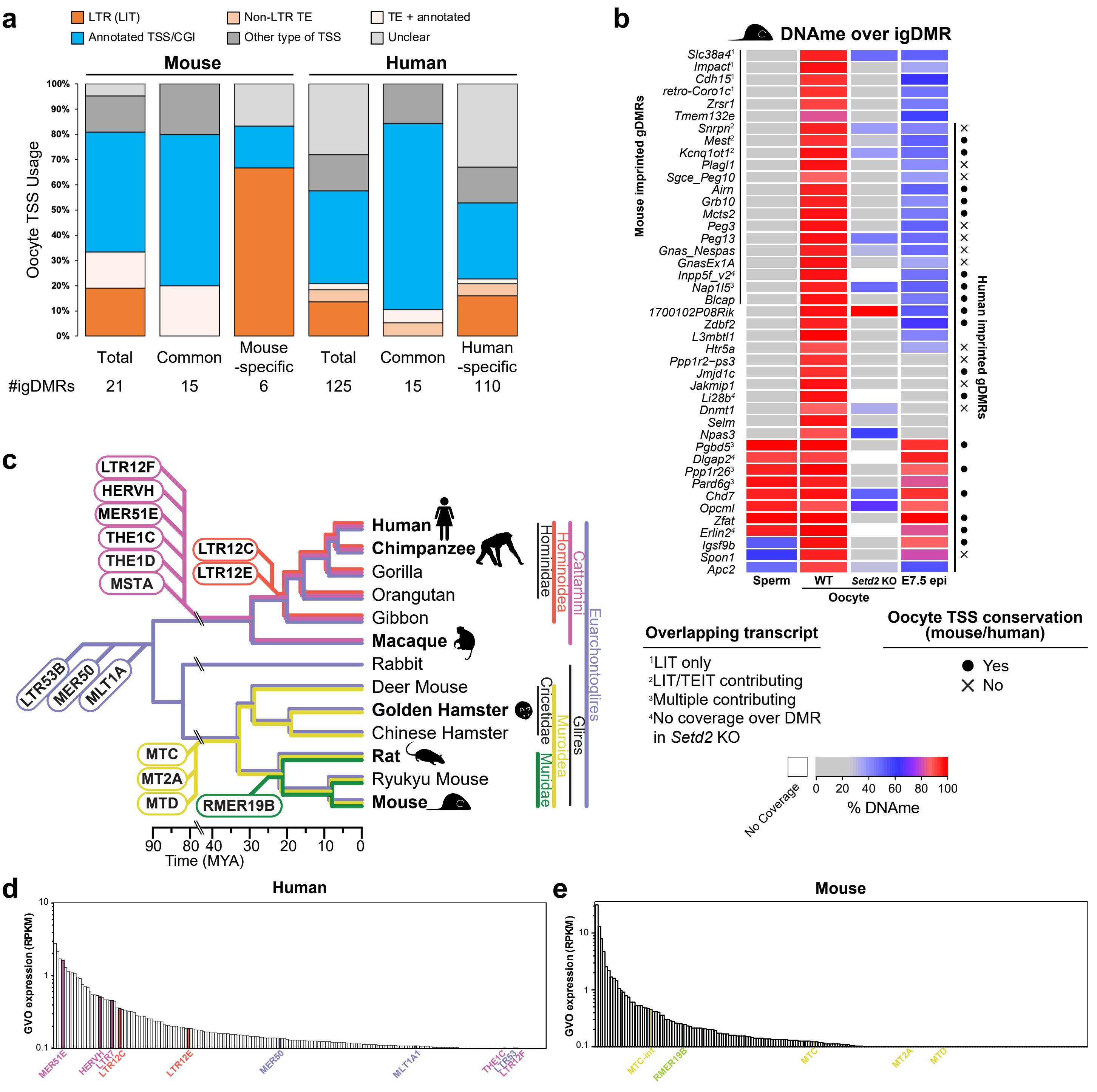


**Supplementary Figure 1. TSS usage, DNAme at igDMRs, LTR colonization time and oocyte expression of LTR families. a.** Percentage of categories of transcription start sites (TSSs) for all oocyte transcripts overlapping the mouse or human igDMRs characterized in this study. TE: transposable element. **b.** Heatmap showing CpG DNAme percentages over 20 mouse igDMRs (black line on left) and mouse regions syntenic with human igDMRs (black line on right), in mouse: mature sperm, wild-type oocytes (WT), *Setd2*-deficient oocytes (*Setd2* KO), and wild-type E7.5 epiblast (E7.5 epi). Categories of overlapping transcripts are identified by superscript numerals and the conservation of the mouse and human oocyte TSS for transcripts overlapping these syntenic regions is highlighted on the right. **c.** Evolutionary colonization time of the LTR families under study, in relation to the phylogeny of catarrhine primates, rodents and glires. Note that among the primate-specific LTR families, LTR12C and LTR12E (red) colonized the common ancestor of modern apes and are therefore absent from the genomes of Old World monkeys, including macaques. Conversely, MTC, MT2A, MTD and RMER19B LTR families colonized the rodent lineage and are therefore absent from rabbit and primates. **d-e.** Agglomerated expression levels (RPKM) of all LTR families (> 100 copies) in human (**d**) and mouse (**e**) oocytes. Colonization time of the families apparently responsible for DNAme over igDMRs color coded as presented in **c**.

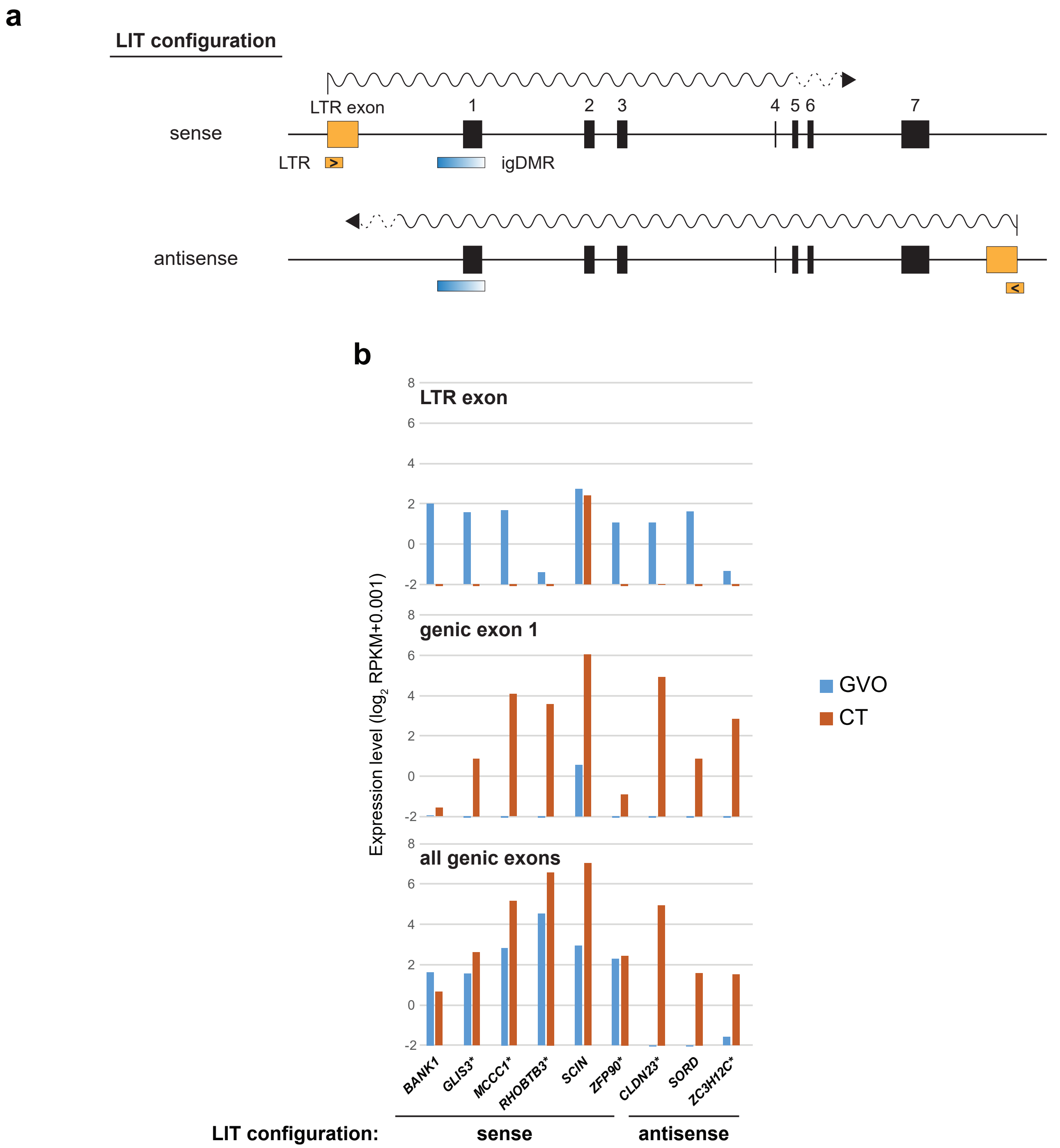


**Supplementary Figure 2. LIT configurations and promoter usage at human igDMRs. a.** Diagram showing the genomic structure of a hypothetical 7-exon gene (top) with a promoter igDMR, and a LIT originating either upstream and sense to the gene, or downstream and antisense to the gene. The LIT configuration for each of the 17 human-specific igDMRs is shown in **Supplementary Table 3**. **b.** RNAseq coverage in purified cytotrophoblast (CT) and oocytes (GVO) for the LTR exon, annotated genic exon 1, and the entire gene (exons 1-n), for the 9 human-specific igDMRs expressed (>0.75 RPKM) in CT. The gene/LIT pairs were grouped based on their respective LIT configuration. The 6 genes which show clear paternal-biased expression in (>70%) CT (**Supplementary Table 3**) are labelled with an asterisk.

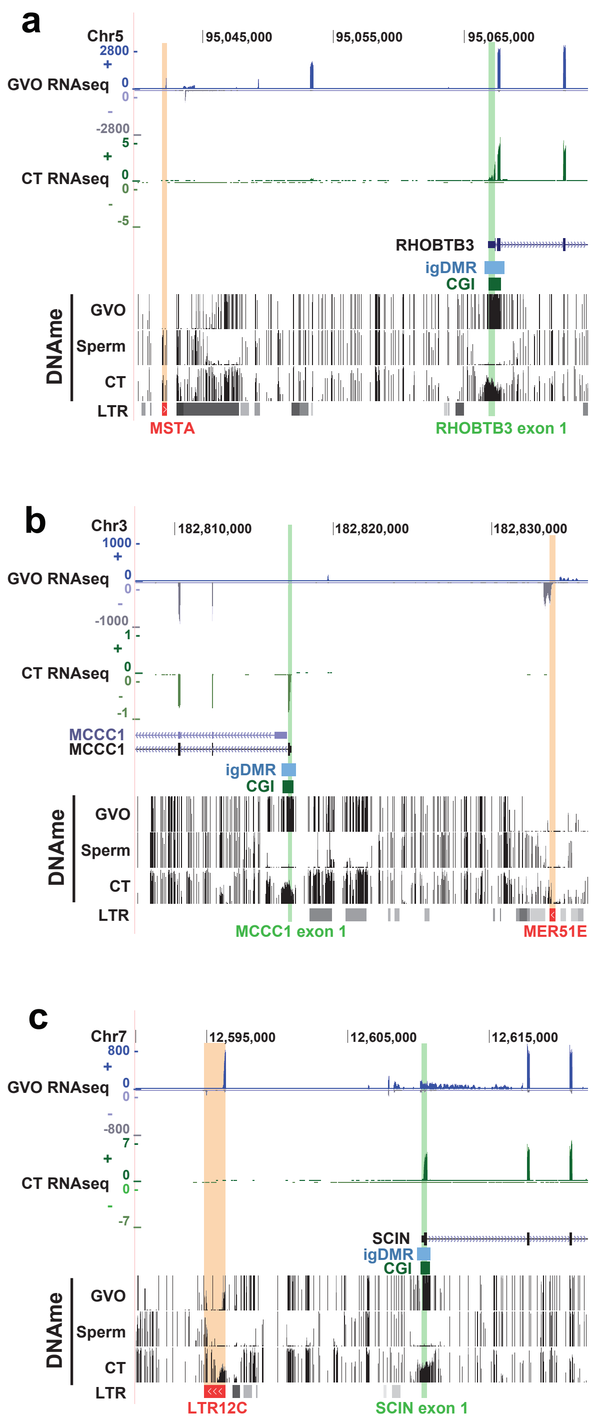


**Supplementary Figure 3. Transcription initiation and DNAme in human oocytes and cytotrophoblast at *RHOBTB3*, *MCCC1*, and *SCIN* loci.** For the igDMRs at *RHOBTB3* (**a**), *MCCC1* (**b**), and *SCIN* (**c**), RNA-seq and DNAme tracks in oocytes (GVO) and purified cytotrophoblast (CT) are shown together with annotations, including promoter exons (LTR exon, canonical exon 1), the igDMR, CGI, and LTR retrotransposons from Repeat Masker. The LTR within which the observed LIT initiates and its orientation are highlighted in red. For both cell types, stranded RNA-seq data are presented separately for the positive (+) and negative (-) strand. Sperm DNAme is also shown to highlight the boundaries of the gDMR in mature gametes.

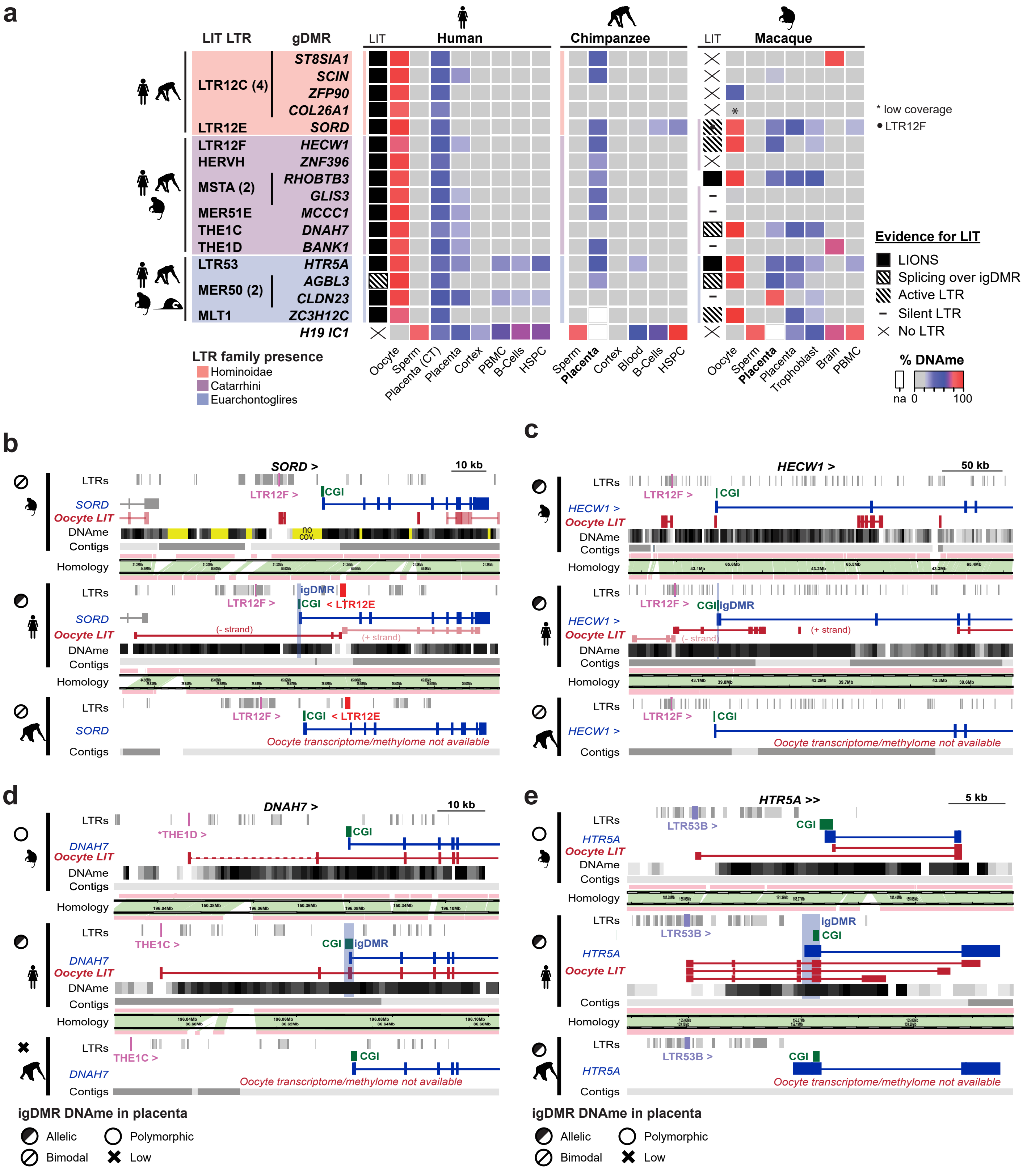


**Supplementary Figure 4. DNA methylation at LIT-associated gDMRs in primates. a.** List of human imprinted placental genes/igDMRs associated with each of the LITs/LTRs, color-coded according to the phylogenetic distribution of the ERV family: colonization of the common ancestor of apes (Hominoidea) in pink, apes and Old World monkeys (Catarrhini) in purple or primates and rodents (Euarchontoglires) in blue. The confirmed or predicted presence of a LIT overlapping with or proximal to the cognate gDMR in humans and macaques is indicated (see methods). The DNAme level (heat map) for each putative igDMR in syntenic regions in the gametes, placenta, brain (cortex) and blood (PBMC, B-cells and HSPC) is also shown (where available) for human, chimpanzee and macaque, as determined from published WGBS datasets. Note that the putative *SORD* igDMR gains *de novo* DNAme as the apparent result of LTR12E- and LTR12F-initiated transcripts in human and macaque oocytes, respectively. Placenta: average DNAme using our targeted bisulfite sequencing data from chimpanzee and macaque. **b-e.** Screenshots of the human *SORD*, *HECW1*, *DNAH7* and *HTR5A* loci and syntenic regions in chimp and macaque. The annotated gene (blue), CGI (green) and human igDMR (blue shading) are shown, as well as oocyte transcripts inferred from *de novo* transcriptome assembly (red) and DNAme (grayscale) in human and macaque oocytes. The relevant LTR, colored by common ancestor (red: Hominoidea, purple: Catarrhini, Euarchontoglires: blue), is highlighted in each species when present. Regions of homology and contigs are also shown. Note that transcriptome and methylome data for chimp oocytes are not available.

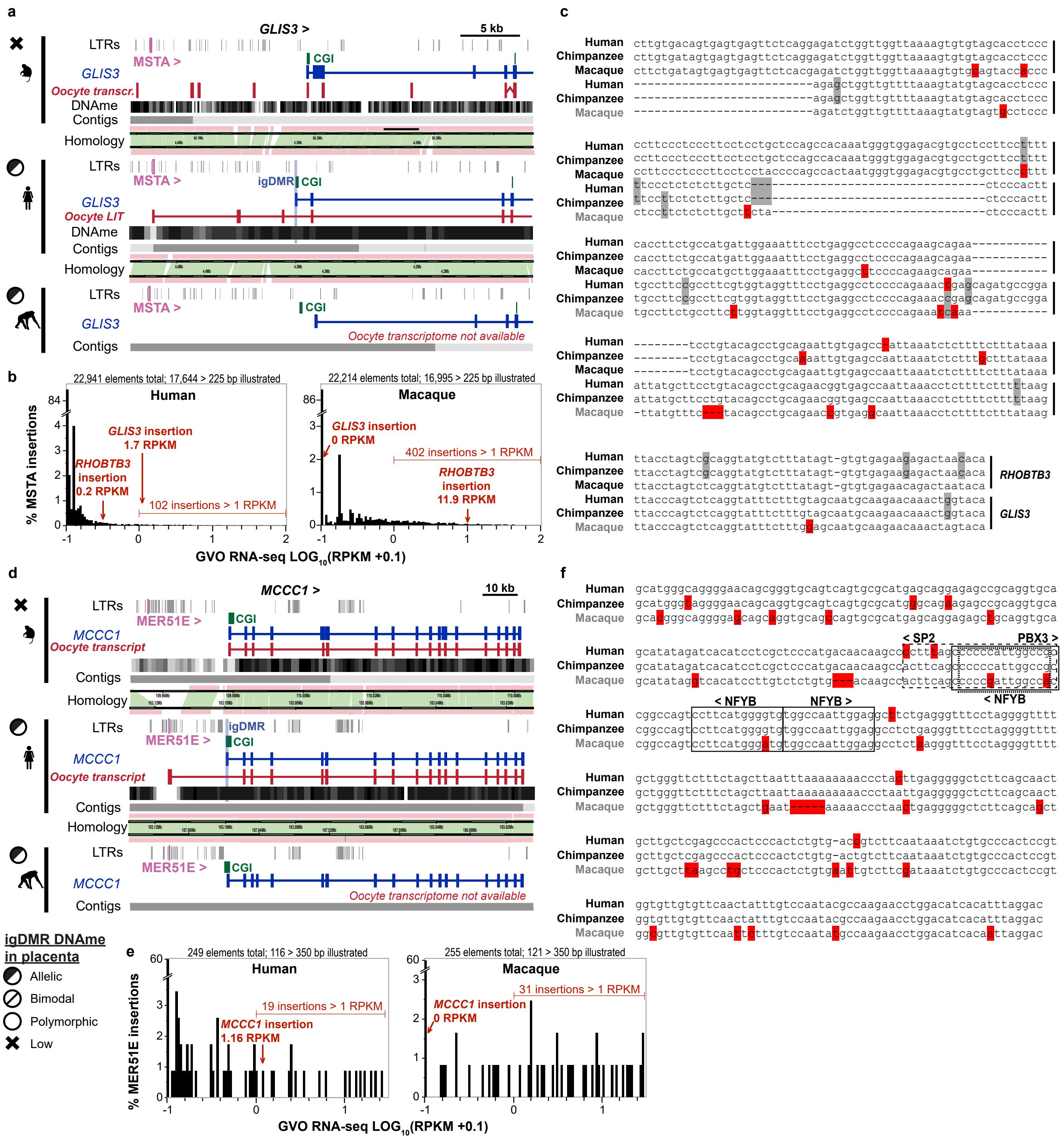


**Supplementary Figure 5. Analyses of MSTA and MER51 elements. a.** Screenshot of the *GLIS3* locus in macaque, human and chimp. The annotated gene (blue), CGI (green) and human gDMR (blue shading) are shown, as well as oocyte transcripts inferred from *de novo* transcriptome assembly (red) and DNAme (grayscale) in human and macaque oocytes. While the gDMR at the 5' end of *GLIS3* is embedded within a LIT initiating in an upstream active MSTA LTR (highlighted in purple) in human oocytes, no LIT is detected in the orthologous region in the macaque locus nor is any RNA-seq coverage detected over the MSTA itself. **b.** Expression levels (RNA-seq coverage in RPKM) in GVOs of all 22,941 and 22,214 MSTA elements (>225 bp) annotated in the human and macaque genomes, respectively. Unique coverage over relevant LTRs upstream of the *RHOBTB3* and *GLIS3* genes is shown for both species. Note that the vast majority of MSTA elements are transcriptionally inert in both species, with greater than 84% and 86% of elements in human and macaque respectively with RPKM=0. The relevant LTR upstream of the *RHOBTB3* gene are expressed in both species, while the MSTA upstream of the *GLIS3* gene is expressed in human (RPKM= 1.7) but not in macaque (RPKM= 0.0). The expression level of these MSTA insertions in chimpanzee oocytes is unknown. **c.** Multiple alignment analysis of the MSTA elements situated upstream of *RHOBTB3* and *GLIS3* in human, chimpanzee and macaque. Polymorphic regions found in a single species and insertion are highlighted in red, and those found in two are highlighted in gray. A number of small deletions and single nucleotide polymorphisms may impact transcription of the macaque LTR. **d.** Screenshot, as in **a**, of the *MCCC1* locus in macaque, human and chimp. Despite the presence of a MER51E (highlighted in purple) upstream of the *MCCC1* gene, no transcript initiating in this LTR is detected in macaque oocytes and no RNA-seq coverage is detected over the element itself. **e.** Expression levels in GVOs of all 249 and 255 MER51 elements (>350 bp) annotated in the human and macaque genomes, respectively. Unique coverage over the relevant LTR upstream of the *MCCC1* gene is shown for both species. Note that the vast majority of MER51 elements are transcriptionally inert in both species and the relevant LTR upstream of the *MCCC1* gene is expressed in human (RPKM= 1.2) but not in macaque oocytes (RPKM= 0.0). **f.** Multiple alignment analysis of the MER51E element situated upstream of *MCCC1* in human, chimpanzee and macaque. Polymorphic regions found in a single species and insertion are highlighted in red, and those found in two are highlighted in gray. Binding motifs for the transcription factors SP2, NFYB and PBX3, which are all expressed in oocytes, have been identified in human[1]. Note the presence of multiple single nucleotide polymorphisms over these transcription factor binding sites between the human and macaque insertions.

1.Ito, J., Sugimoto, R., Nakaoka, H., Yamada, S., Kimura, T., Hayano, T. & Inoue, I. Systematic identification and characterization of regulatory elements derived from human endogenous retroviruses. PLoS Genet. 13, e1006883 (2017).

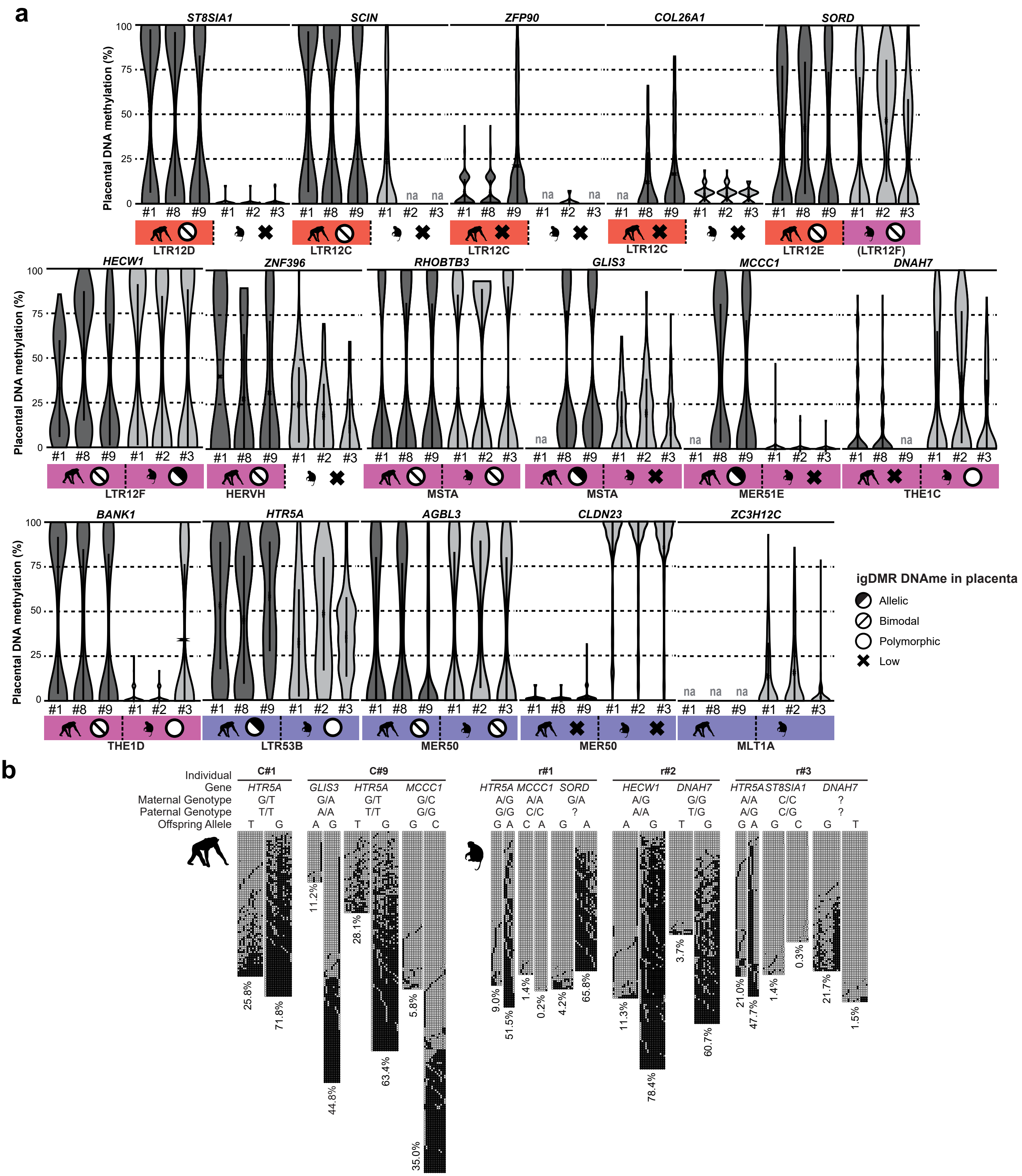


**Supplementary Figure 6. Targeted bisulfite analysis of LIT-associated gDMRs in chimp and macaque placenta. a.** Violin plots showing the distribution of mean DNAme levels of individual bisulphite-sequencing reads covering the regions syntenic to the gDMRs of selected genes in placentae from chimp (individuals 1, 8, and 9) and macaque (individuals 1, 2, and 3). The presence of the cognate proximal LTR is shown below, color coded as in Figure 1b. na= data not available. **b.** DNAme data (each row represents a sequenced molecule) are shown for genes and individual placentae for which allelic data was available. The individual chimp or rhesus macaque placenta from which DNA was extracted is also shown along with maternal and paternal genotype at the informative SNP, when available. Filled and open circles represent methylated and unmethylated CpGs, respectively. Average percent methylation for parental strands is shown at the bottom.

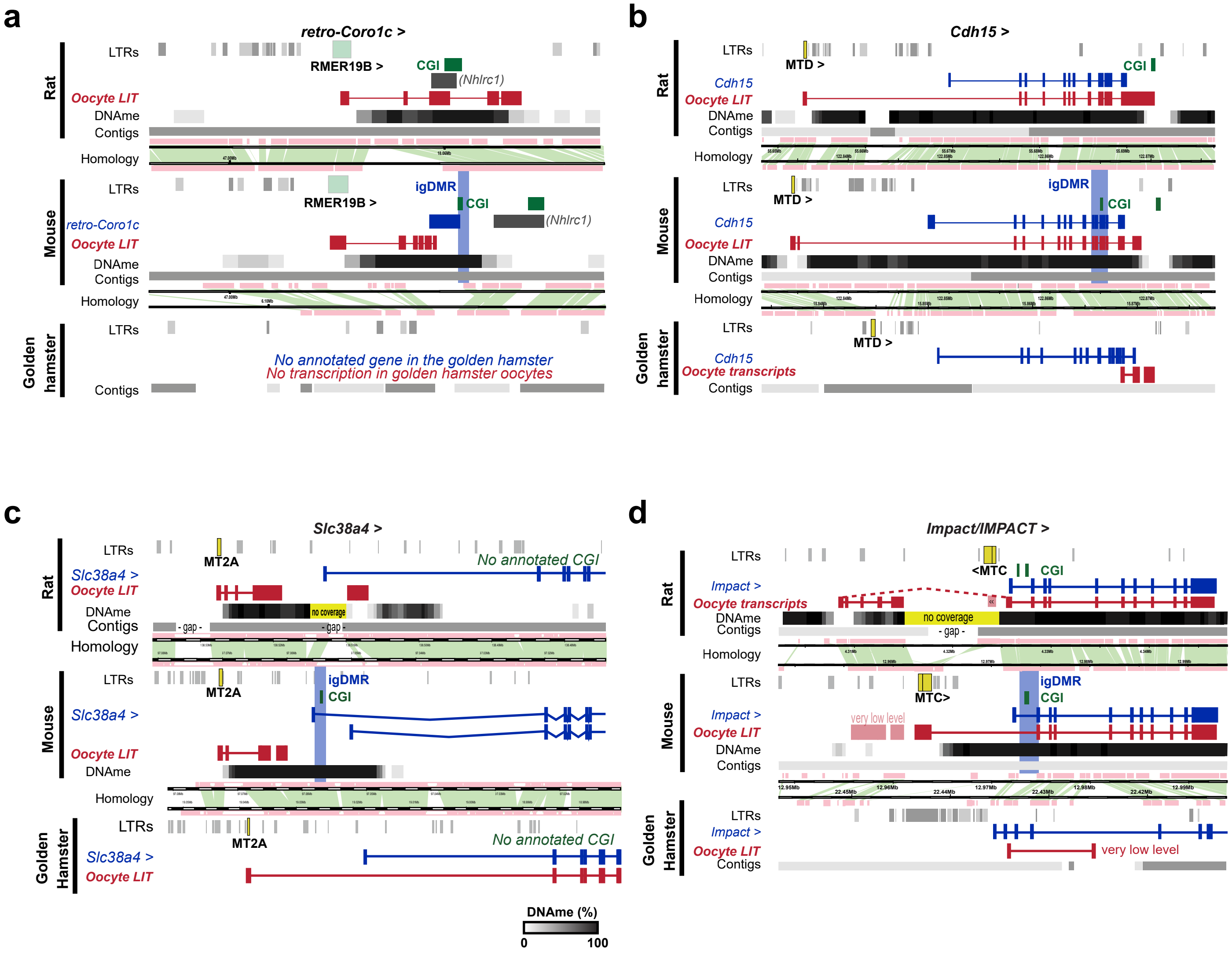


**Supplementary Figure 7. Mouse-specific igDMRs.** Screenshots of the mouse *retro-Coro1c* (**a**), *Cdh15* (**b**), *Slc38a4* (**c**), and *Impact* (**d**) loci, and syntenic regions in rat and golden hamster. The annotated gene (blue), CGI (green) and human igDMR (blue shading) are shown, as well as oocyte transcripts inferred from *de novo* transcriptome assembly. The upstream LTRs (green/yellow) acting as oocyte promoters for the LITs are highlighted in all three species, and downstream DNAme blocks are shown (mouse and rat oocytes; grayscale). Regions of homology and genomic contigs are also shown.

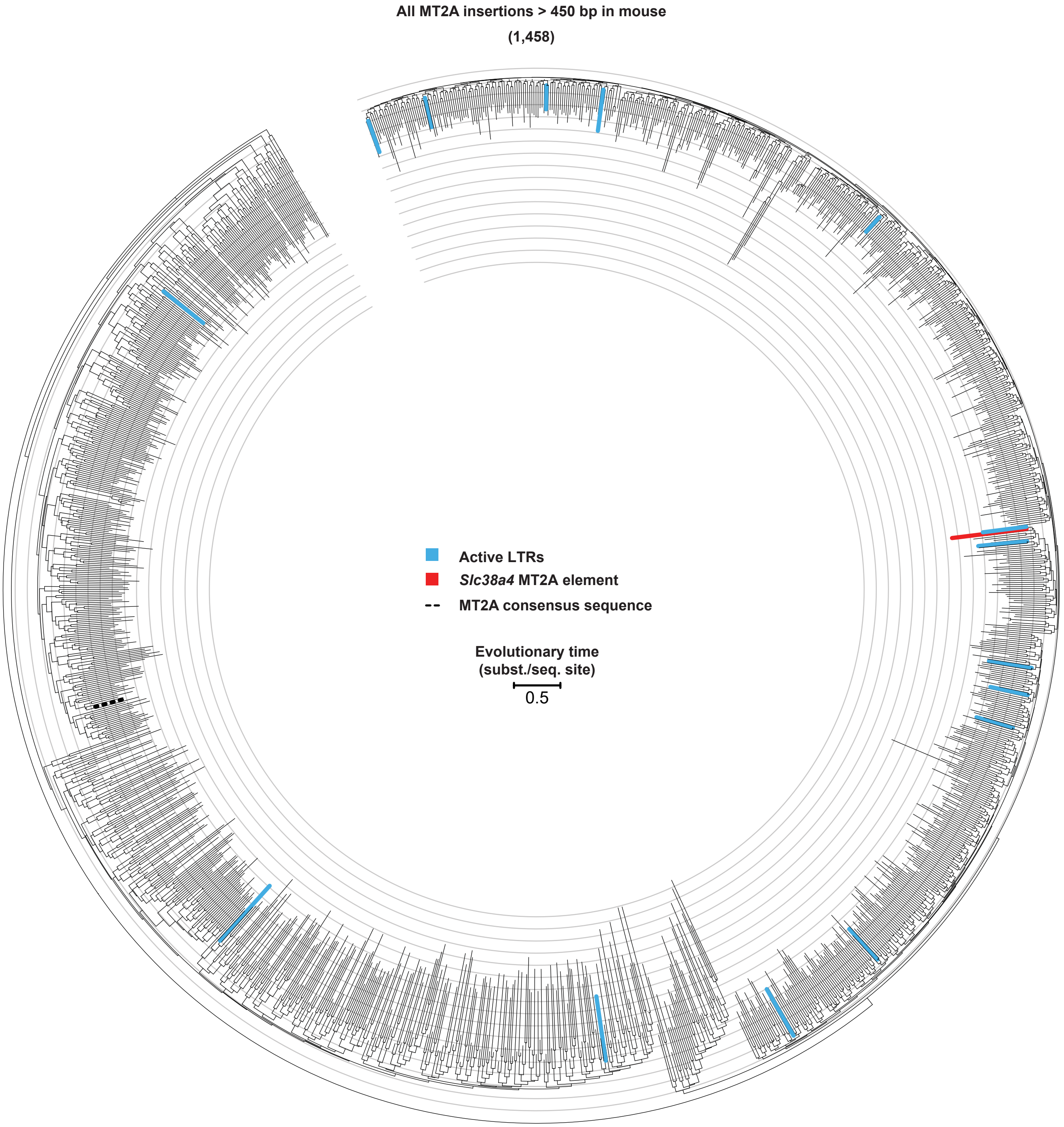


**Supplementary Figure 8. MT2A evolutionary tree.** Phylogenetic tree illustrating 1,458 annotated MT2A elements > 450 bp (RepeatMasker). MT2A LTRs in which transcript initiation is detected in oocytes based on uniquely aligned reads (RNAseq) are highlighted in blue. The MT2A LTR promoter driving the LIT upstream of the *Slc38a4* igDMR is highlighted in red, and the consensus MT2A sequence is highlighted in black dashes.

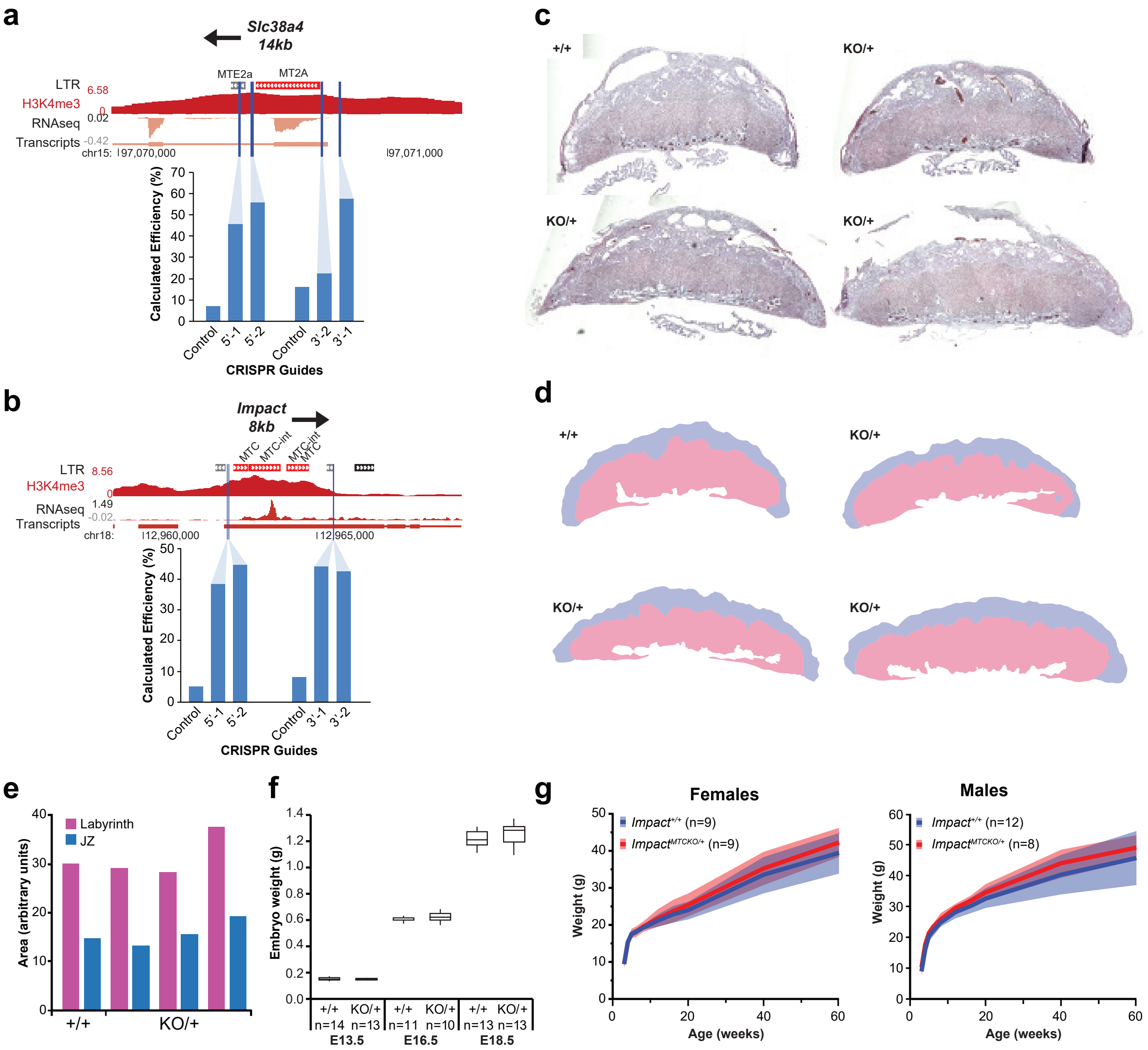


**Supplementary Figure 9. CRISPR-Cas9 deletion strategy and phenotype analysis. a-b.** Screenshots of the (**a**) MT2A LTR initiating a LIT upstream of *Slc38a4*, and (**b**) the MTC element initiating a LIT upstream of *Impact* in mouse. H3K4me3 and RNA-seq from GVO are shown, as well as oocyte transcripts inferred from *de novo* transcriptome assembly. The transcriptional orientation of both genes and the distance to their respective annotated canonical exon 1 is shown above. The locations of protospacers for the CRISPR-Cas9 gRNAs tested are highlighted by blue lines, and the calculated mutagenic efficiency of each guide, as determined by T4 Endonuclease I assay in C2 ESCs, is shown below. **c.** H&E staining of E13.5 placentae from three *Slc38a4*$^{MT2AKO/+}$ conceptuses and a wild-type littermate. **d.** Highlighted labyrinth layer (pink) and junctional zone (JZ, blue) of *Slc38a4*$^{MT2AKO/+}$ and wild-type E13.5 placentae. **e.** Quantitation of relative area of labyrinth and JZ from *Slc38a4*$^{MT2AKO/+}$ and wild-type E13.5 placentae. Each set of bars represents an individual sample. **f.** Weight of *Slc38a4*$^{MT2AKO/+}$ embryos and wild-type littermates at E13.5, E16.5, and E18.5. The number of embryos weighed is indicated underneath the genotypes. Box: 25th, 50th and 75th percentile, whiskers: minimum and maximum. **g.** Growth curve of the wild-type and mutant progeny from *Impact*$^{+/MTCKO}$ females crossed to wild-type males. Females (left) and males (right) were weighed at postnatal weeks 3, 4, 5, 6, 8, 12, 16, 20, 40, and 60. No significant weight difference was observed between maternal MTCKO and wild-type littermates. Graphs show mean (bold line) ± standard deviation (shaded area). The number of animals weighed is shown next to the genotypes.

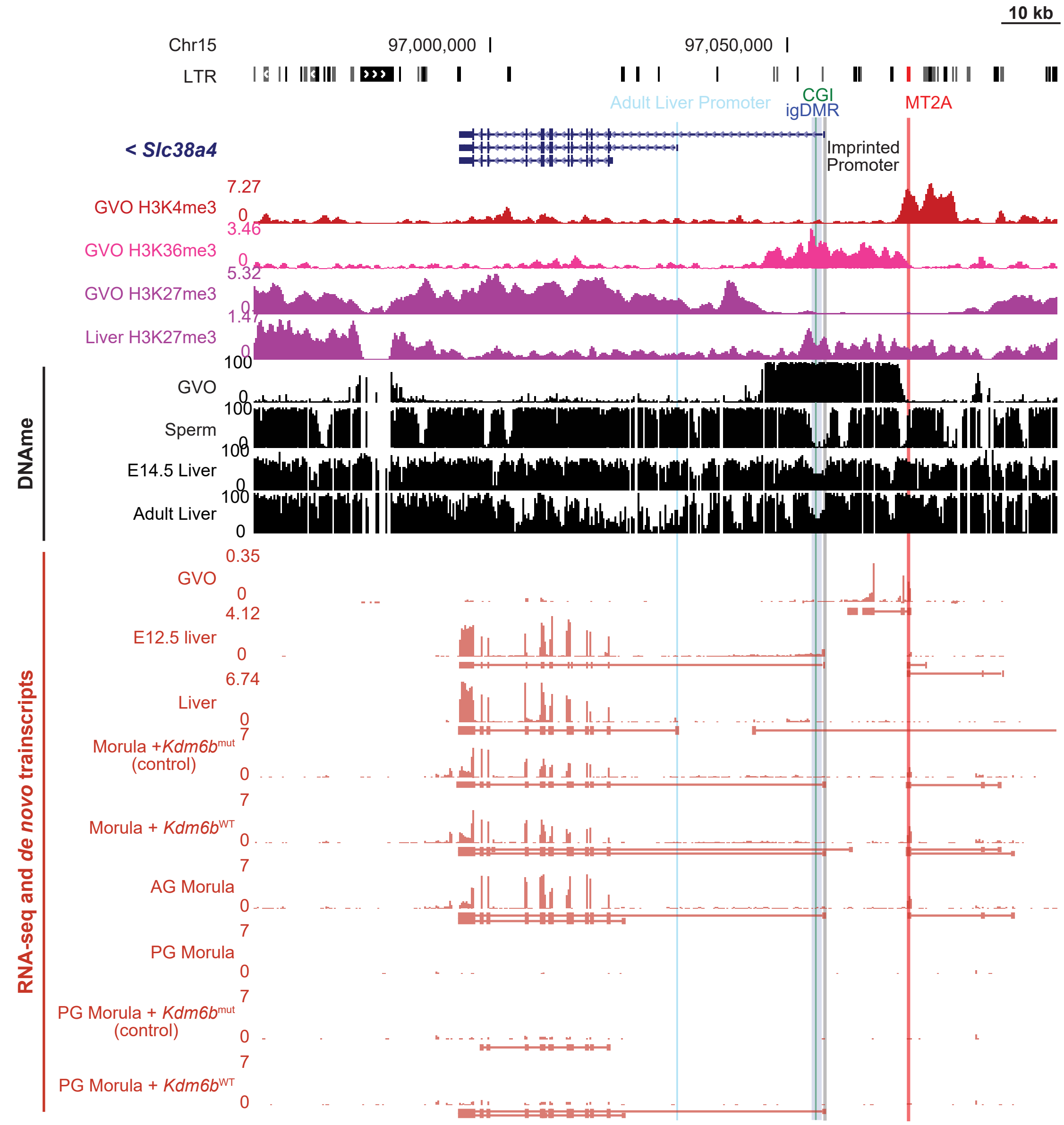


**Supplementary Fig. 10. Promoter usage and epigenetic features at the *Slc38a4* locus.** Screenshot of the *Slc38a4* locus, showing ChIP-seq data, DNAme and *de novo* transcriptome assembly in various developmental stages and tissues. Annotated RefSeq transcript isoforms are shown on top, along with the CGI (green), igDMR (blue), and MT2A element (red) in which the relevant oocyte LIT initiates upstream of the *Slc38a4* gene. H3K4me3 enrichment over the MT2A element in GVO is shown in red. Mutually exclusive H3K36me3 and H3K27me3 marks in GVO are shown below, and the loss of the H3K27me3 domain over the gene body of *Slc38a4* in adult liver is shown underneath. DNAme from mature gametes, developing and adult liver, as well as IVF and SCNT morulae are shown in black. RNA-seq from developing (E12.5) and adult liver shows the switch in promoter usage, in parallel with the observed switch from a paternal (imprinted) to biallelic transcription pattern. RNA-seq profiles from biparental or parthenogenetic (PG) morulae microinjected with mutant (control) or wild-type *Kdm6b* transcripts (H3K27 demethylase) are from Inoue et al. 2017[1]. While injection of *Kdm6b* in 1-cell embryos was reported to yield an increase in maternal *Slc38a4* transcripts at the morula stage, it is not possible to determine whether these low-level maternal transcripts initiate from the imprinted (igDMR) or the biallelic adult liver promoter (highlighted in gray and light blue, respectively). Moreover, *Kdm6b* injection in PG 1-cell embryos does not recapitulate the transcription levels observed in biparental control morulae.

1. Inoue, A., Jiang, L., Lu, F., Suzuki, T. & Zhang, Y. Maternal H3K27me3 controls DNA methylation-independent imprinting. Nature 547, 419–424 (2017).

**Supplementary Table 1. Datasets used in this study**

| Species | Tissue | Data type | Accession | Reference |
|---|---|---|---|---|
| Human | F-PGC 10,11W | WGBS | GSE63818 | Guo, F. et al. Cell 161, 1437-52 (2015). |
| Human | Oocyte | WGBS | DRA003802 | Okae, H. et al. PLoS Genet. 10, e1004868 (2014). |
| Human | Oocyte | RNA-seq | GSE85632 | Hendrickson, P. G. et al. Nat. Genet. 487, 57 (2017). |
| Human | Blastocyst | WGBS | DRA003802 | Okae, H. et al. PLoS Genet. 10, e1004868 (2014). |
| Human | Cortex | WGBS | GSE25930 | Schroeder, D. I., Lott, P., Korf, I. & LaSalle, J. M. Genome Res. 21, 1583–1591 (2011) |
| Human | Cytotrophoblast | WGBS | JGA00000000074 | Okae, H. et al. Cell Stem Cell 22, 50–63.e6 (2018). |
| Human | Liver | WGBS | GSM916049 | Roadmap Epigenomics Consortium et al. Nature 518, 317–330 (2015) |
| Human | Lung | WGBS | GSM983647 | Roadmap Epigenomics Consortium et al. Nature 518, 317–330 (2015) |
| Human | Placenta | WGBS | GSM1186665 | Roadmap Epigenomics Consortium et al. Nature 518, 317–330 (2015) |
| Human | Sperm | WGBS | DRA003802 | Okae, H. et al. Cell Stem Cell 22, 50–63.e6 (2018). |
| Human | Thymus | WGBS | GSM1010979 | Roadmap Epigenomics Consortium et al. Nature 518, 317–330 (2015) |
| Mouse | Cortex | WGBS | GSE42836 | Hon, G. C. et al. Nat. Genet. 45, 1198–1206 (2013). |
| Mouse | Oocyte | H3K4me3 ChIPseq | GSE93941 | Hanna, C. W. et al. Nat Struct Mol Biol 25, 73–82 (2018) |
| Mouse | Oocyte | H3K36me3 ChIPseq | GSE112622 | Brind'Amour, J. et al. Nature Communications 9, 3331–3344 (2018) |
| Mouse | Oocyte | PolII ChIPseq | GSE126363 | **This study** |
| Mouse | Liver | H3K27me3 ChIPseq | GSE113326 | Lee, Y.-Y. et al. Nucleic Acids Res 46, 8832–8847 (2018) |
| Mouse | Oocyte | RNA-seq | GSE112622 | Brind'Amour, J. et al. Nature Communications 9, 3331–3344 (2018) |
| Mouse | Morula (Kdm6b Mut injection) | RNA-seq | GSE92605 | Inoue, A., Jiang, L., Lu, F., Suzuki, T. & Zhang, Y. Nature 547, 419–424 (2017) |
| Mouse | Morula (Kdm6b WT injection) | RNA-seq | GSE92605 | Inoue, A., Jiang, L., Lu, F., Suzuki, T. & Zhang, Y. Nature 547, 419–424 (2017) |
| Mouse | Morula (parthenogenetic, Kdm6b Mut injection) | RNA-seq | GSE92605 | Inoue, A., Jiang, L., Lu, F., Suzuki, T. & Zhang, Y. Nature 547, 419–424 (2017) |
| Mouse | Morula (parthenogenetic, Kdm6b WT injection) | RNA-seq | GSE92605 | Inoue, A., Jiang, L., Lu, F., Suzuki, T. & Zhang, Y. Nature 547, 419–424 (2017) |
| Mouse | Morula (androgenetic) | RNA-seq | GSE92605 | Inoue, A., Jiang, L., Lu, F., Suzuki, T. & Zhang, Y. Nature 547, 419–424 (2017) |
| Mouse | Morula (parthenogenetic) | RNA-seq | GSE92605 | Inoue, A., Jiang, L., Lu, F., Suzuki, T. & Zhang, Y. Nature 547, 419–424 (2017) |
| Mouse | Blastocyst (IVF) | RNA-seq | GSE112546 | Matoba, S. et al. Cell Stem Cell. 23, 343-354.e5 (2018). |
| Mouse | Blastocyst (SCNT) | RNA-seq | GSE112546 | Matoba, S. et al. Cell Stem Cell. 23, 343-354.e5 (2018). |
| Mouse | E12.5 Embryo | RNA-seq | GSE82853/4 | ENCODE Project Consortium. Nature 489, 57-74 (2012). |
| Mouse | Oocyte | WGBS | DRA000570 | Shirane, K. et al. PLoS Genet. 9, e1003439 (2013). |
| Mouse | Blastocyst | WGBS | DRA006680 | Brind'Amour, J. et al. Nature Communications 9, 3331–3344 (2018) |
| Mouse | IVF Blastocyst | WGBS | GSE112528 | Matoba, S. et al. Cell Stem Cell 23, 343-354.e5 (2018). |
| Mouse | SCNT Blastocyst | WGBS | GSE112528 | Matoba, S. et al. Cell Stem Cell 23, 343-354.e5 (2018). |
| Mouse | E10.5 embryo | WGBS | GSE58108 | Harten, S.K. et al. BMC Biol. 13:21 (2015). |
| Mouse | E14.5 liver | WGBS | GSE58108 | Harten, S.K. et al. BMC Biol. 13:21 (2015). |
| Mouse | Liver (young) | WGBS | GSE92486 | Hahn, O. et al. Genome Biol. 18(1):56 (2017). |
| Mouse | Liver (aged) | WGBS | GSE92486 | Hahn, O. et al. Genome Biol. 18(1):56 (2017). |
| Mouse | Liver | WGBS | GSE42836 | Hon, G. C. et al. Nat. Genet. 45, 1198–1206 (2013). |
| Mouse | Lung | WGBS | GSE42836 | Hon, G. C. et al. Nat. Genet. 45, 1198–1206 (2013). |
| Mouse | Pancreas | WGBS | GSE42836 | Hon, G. C. et al. Nat. Genet. 45, 1198–1206 (2013). |
| Mouse | Placenta | WGBS | GSE42836 | Hon, G. C. et al. Nat. Genet. 45, 1198–1206 (2013). |
| Mouse | Sperm | WGBS | DRA000484 | Kobayashi, H. et al. PLoS Genet. 8, e1002440 (2012). |
| Mouse | Thymus | WGBS | GSE42836 | Hon, G. C. et al. Nat. Genet. 45, 1198–1206 (2013). |
| Chimpanzee | Sperm | WGBS | GSE30340 | Molaro, A. et al. Cell 146, 1029–1041 (2011) |
| Chimpanzee | Cortex | WGBS | GSE37202 | Zeng, J. et al. Am J Hum Genet 91, 455–465 (2012). |
| Chimpanzee | B-cells | WGBS | SRP059313 | Hodges, E. et al. Mol Cell 44, 17–28 (2011). |
| Chimpanzee | Blood | WGBS | SRP059313 | Hernando-Herraez, I. et al. Nucleic Acids Res 43, 8204–8214 (2015) |
| Chimpanzee | HSPC | WGBS | SRP059313 | Hodges, E. et al. Mol Cell 44, 17–28 (2011). |
| Rhesus Macaque | Oocyte | RNA-seq | GSE112534 | Ruebel, M. L. et al. Molecular Human Reproduction 24, 478–494 (2018) |
| Rhesus Macaque | Oocyte | WGBS | GSE60166 | Gao, F. et al. Cell Res 27, 526–539 (2017). |
| Rhesus Macaque | Sperm | WGBS | GSE60166 | Gao, F. et al. Cell Res 27, 526–539 (2017). |
| Rhesus Macaque | Placenta | WGBS | GSE63330 | Schroeder, D. I. et al. PLoS Genet. 11, e1005442 (2015). |
| Rhesus Macaque | Trophoblast | WGBS | GSE63330 | Schroeder, D. I. et al. PLoS Genet. 11, e1005442 (2015). |
| Rhesus Macaque | Brain | WGBS | GSE77124 | Mendizabal, I. et al. Molecular Biology and Evolution 33, 2947–2959 (2016) |
| Rhesus Macaque | PBMC | WGBS | GSE34128 | Tung, J. et al. Proc Natl Acad Sci USA 109, 6490–6495 (2012) |
| Rat | Oocyte | RNA-seq | GSE112622 | Brind'Amour, J. et al. Nature Communications 9, 3331–3344 (2018) |
| Rat | Oocyte | WGBS | DRA006680 | Brind'Amour, J. et al. Nature Communications 9, 3331–3344 (2018) |
| Golden Hamster | Oocyte | RNA-seq | GSE86470 | Franke, V. et al. Genome Res. 27, 1384–1394 (2017). |

**Supplementary Table 3: Transcription initiation related to LIT-associated igDMRs in oocytes and first trimester cytotrophoblasts**

| | | | | | | LIT configuration | | | | | oocyte expression | | CT expression | | | |
|---|---|---|---|---|---|---|---|---|---|---|---|---|---|---|---|---|
| **igDMR hg19 coordinates** | **CGI?** | **Human Gene** | **Gene orientation** | **LTR** | **LTR hg19 coordinates** | **LTR orientation** | **LTR position relative to gene** | **LIT orientation** | **LIT orientation vs gene** | **Chimeric transcript in oocyte** | **main oocyte TSS** | **oocyte, gene RPKM** | **main CT TSS** | **CT TSS overlap with igDMR** | *** % paternal allele expression** | **CT, gene RPKM** |
| chr4:102711702-102712397 | *Yes* | *BANK1* | + | THE1D | chr4:102386486-102386876 | + | 5' | + | S | **yes** | LTR | 3.128 | Annotated Exon 1 | **yes** | - | 0.903 |
| chr7:101006052-101006963 | *Yes* | *COL26A1* | + | LTR12C | chr7:100991675-100993150 | + | 5' | + | S | **yes** | LTR | 0.267 | Annotated Exon 1 (very low) | **yes** | - | 0.325 |
| chr2:196933266-196934154 | *Yes* | *DNAH7* | – | THE1C | chr2:196972811-196973166 | – | 5' | – | S | **yes** | LTR | 0.826 | Not expressed | **-** | - | 0.002 |
| chr9:4298157-4300432 | *Yes* | *GLIS3* | – | MSTA | chr9:4419277-4419507 | – | 5' | – | S | **yes** | LTR | 2.947 | Annotated Exon 1 | **yes** | 84.1 | 6.030 |
| chr7:43151828-43153950 | *Yes* | *HECW1* | + | LTR12F | chr7:43116241-43116507 | + | 5' | + | S | **yes** | LTR | 1.071 | Not expressed | **-** | - | 0.052 |
| chr7:154861569-154863381 | *Yes* | *HTR5A* | + | LTR53 | chr7:154851368-154851877 | + | 5' | + | S | **yes** | LTR | 7.339 | Not expressed | **-** | - | 0.000 |
| chr3:182816738-182817626 | *Yes* | *MCCC1* | – | MER51E | chr3:182833657-182834013 | – | 5' | – | S | **yes** | LTR | 7.178 | Annotated Exon 1 | **yes** | 95.0 | 35.290 |
| chr5:95066568-95068092 | *Yes* | *RHOBTB3* | + | MSTA | chr5:95041914-95042302 | + | 5' (*SPATA9* intron) | + | S | **yes** | LTR | 22.817 | Annotated Exon 1 | **yes** | 90.2 | 94.975 |
| chr7:12609907-12610833 | *Yes* | *SCIN* | + | LTR12C | chr7:12594828-12596347 | – | 5' | + | S | **yes** | LTR | 7.639 | Annotated Exon 1 (and LTR contribution) | **yes** | 63.9 | 132.834 |
| chr12:22487219-22488465 | *Yes* | *ST8SIA1* | – | LTR12C | chr12:22529637-22530123 | + | 5' | – | S | **yes** | LTR | 0.310 | Not determined | **-** | - | 0.230 |
| chr7:138348963-138349444 | *Yes* | *SVOPL* | – | MLT1A1 | chr7:138361021-138361358 | – | 5' | + | S | **yes** | LTR | 140.245 | chr7:138342501-138365256 | **no** | 20.5 | 8.490 |
| chr16:68572892-68573971 | *Yes* | *ZFP90* | + | LTR12C | chr16:68560468-68562139 | + | 5' | + | S | **yes** | LTR | 4.934 | Annotated Exon 1 | **yes** | 73.0 | 5.384 |
| chr7:134671024-134671987 | *Yes* | *AGBL3* | + | MER50 | chr7:134762504-134762774 | – | intron | – | AS | no | LTR | 0.032 | Annotated Exon 1 (very low) | **yes** | - | 0.053 |
| chr8:8559131-8560867 | *Yes* | *CLDN23* | + | MER50 | chr8:8639704-8640372 | + | 3' | – | AS | no | LTR | 0.027 | Annotated Exon 1 (monoexonic) | **yes** | 76.1 | 30.647 |
| chr15:45314789-45315642 | *Yes* | *SORD* | + | LTR12E | chr15:45326850-45328464 | + | intron | – | AS | no | LTR | 0.088 | Annotated exon 1 | **yes** | - | 2.979 |
| chr11:109963338-109964976 | *Yes* | *ZC3H12C* | + | MLT1A0 | chr11:109990326-109990684 | – | intron | – | AS | no | LTR | 0.334 | chr11:109963826-109963950 | **yes** | 95.0 | 2.506 |
| chr18:32956850-32957683 | *Yes* | *ZNF396* | – | HERVH | chr18:32938607-32941345 | – | 3' | + | AS | no | LTR | 0.014 | Annotated exon 1 | **yes** | - | 0.250 |

*As reported by Hamada (2016).
Hamada, H., Okae, H., Toh, H., Chiba, H., Hiura, H., Shirane, K., Sato, T., Suyama, M., Yaegashi, N., Sasaki, H. & Arima, T. Allele-Specific Methylome and Transcriptome Analysis Reveals Widespread Imprinting in the Human Placenta. Am. J. Hum. Genet. 99,

**Supplementary Table 4. Oligonucleotide primers used in this study**

| Gene | Species | Name | Sequence (5' to 3') |
|---|---|---|---|
| – | – | XbaI-R gRNA | GGCGCACGCACTAAAAAATCGTCTAGAAAAAAAGCACCGACTCGGTGCCACTTTTTCAAGTTGATAACGGACTAGCCTTATTTTAACTTGCTATTTCTAG |
| *Impact* | mouse | SapI-I5-1 | TGTGGAAAGGGCTCTTCACCGacacaaacctatccgaggtcGTTTTAGAGCTAGAAATAGCAAGTTAAAATAAGGCTAG |
| *Impact* | mouse | SapI-I5-2 | TGTGGAAAGGGCTCTTCACCGtatagtgggaagtccctcatGTTTTAGAGCTAGAAATAGCAAGTTAAAATAAGGCTAG |
| *Impact* | mouse | SapI-I3-1 | TGTGGAAAGGGCTCTTCACCGtaagaccttggtcagcatacGTTTTAGAGCTAGAAATAGCAAGTTAAAATAAGGCTAG |
| *Impact* | mouse | SapI-I3-2 | TGTGGAAAGGGCTCTTCACCGaccctccttgtctaagacctGTTTTAGAGCTAGAAATAGCAAGTTAAAATAAGGCTAG |
| *Impact* | mouse | MTC 5' F1 | AGGCCATGGAAATCTCAAAA |
| *Impact* | mouse | MTC 5' R1 | GCATGAGACGATAGAATTGTGC |
| *Impact* | mouse | MTC 3' F1 | AGGGGTGTGGGGATTAAAAG |
| *Impact* | mouse | MTC 3' R1 | AAAGGGGTATGGAGGGACTG |
| *Impact* | mouse | RFLP-IF1 | ATCGAACAGCCTTGAGGAGA |
| *Impact* | mouse | RFLP-IR1 | GGAGGAAACACAATGCTGGT |
| *Impact* | mouse | BS-IC-F2 | TTGTAAGAAATAGAATTTAGGGTAAG |
| *Impact* | mouse | BS-IC-R2 | AAACTATAACCCCAAAATAACCACTAATA |
| *Impact* | mouse | IqF1 | CATTAAGACGTGAACCTGTGTGAGA |
| *Impact* | mouse | IqR1 | AACTGGCTGACTTGAGCTGTCA |
| *Slc38a4* | mouse | SapI-S5-1 | TGTGGAAAGGGCTCTTCACCGtcttacttgaagacgcatccGTTTTAGAGCTAGAAATAGCAAGTTAAAATAAGGCTAG |
| *Slc38a4* | mouse | SapI-S5-2 | TGTGGAAAGGGCTCTTCACCGaatgagcaaaccacgtgaaaGTTTTAGAGCTAGAAATAGCAAGTTAAAATAAGGCTAG |
| *Slc38a4* | mouse | SapI-S3-1 | TGTGGAAAGGGCTCTTCACCGtcaagtctttgccgatgtacGTTTTAGAGCTAGAAATAGCAAGTTAAAATAAGGCTAG |
| *Slc38a4* | mouse | SapI-S3-2 | TGTGGAAAGGGCTCTTCACCggtggcttaggacagtatcgGTTTTAGAGCTAGAAATAGCAAGTTAAAATAAGGCTAG |
| *Slc38a4* | mouse | MT2A F1 | CTGTGAAAACCATCCCATCC |
| *Slc38a4* | mouse | MT2A R1 | CTACTCGGCTGCTGAGGAAC |
| *Slc38a4* | mouse | MT2A F2 | TGGGTCTCTCCACATCCAAT |
| *Slc38a4* | mouse | MT2A R2 | GGCCTGTTTGAACCAACTGT |
| *Slc38a4* | mouse | RFLP-SF1 | GATTGGGGCTTTGGTCTTCC |
| *Slc38a4* | mouse | RFLP-SR1 | TGTAGAGTGAGAGATGGCCG |
| *Slc38a4* | mouse | SqF1 | ACTGACCGTGCCCATCGT |
| *Slc38a4* | mouse | SqR1 | TTGAGTGCGATGATGATTGCA |
| *Slc38a4* | mouse | BS-SF2 | AGTGTTTTAAGGTAAAAGGAGTT |
| *Slc38a4* | mouse | BS-SR1 | CTCTAAAACTTAAAAACTCTAAAAA |
| *Slc38a4* | mouse | BS-SR2 | CCAAAAACCTAAAACAAAATAAAAAT |
| *AGBL3* | chimp | BS-F | GAGAGTTGGGAGTGTGGTTGG |
| *AGBL3* | chimp | BS-R | CCTTACTTCTACATACCTCACC |
| *AGBL3* | rhesus | BS-F | GTGAGGTAGGTAGAAGGAAGGT |
| *AGBL3* | rhesus | BS-R | TCCCRACAATACCAAAAAATTTTTAAC |
| *BANK1* | chimp | BS-F | AGGTTTTTGAGTAGTTTTATTTTTTTTGGG |
| *BANK1* | chimp | BS-R | ATTTTCTCTTCCTTTAACCCTC |
| *BANK1* | rhesus | BS-F | GGGTAAGGGGTTTGGAAGTT |
| *BANK1* | rhesus | BS-R | ACRCCCTTACCTCAATACCC |
| *CLDN23* | chimp | BS-F | TAGGAGGGAATTAGTTTAAGTGGG |
| *CLDN23* | chimp | BS-R | CCCCTACAACCTACCTTCCTTC |
| *CLDN23* | rhesus | BS-F | GAGGTAGAGTTTTGGGATTGGG |
| *CLDN23* | rhesus | BS-R | CAACACCATCCAAATAAAATAACCC |
| *COL26A1* | chimp | BS-F | GGGATTGGGGGTTGTAGATAT |
| *COL26A1* | chimp | BS-R | CTACACCCAAAACCCTTCAATC |
| *COL26A1* | rhesus | BS-F | GGTGGTTATGGYGTTATTTGT |
| *COL26A1* | rhesus | BS-R | CRCCCAATCTTTAAAACTAAAAC |
| *DNAH7* | chimp | BS-F | GGGTAAATAGTTGGGAAGGGAAT |
| *DNAH7* | chimp | BS-R | AATAAAAAAACTACTCTTTACCAAACTC |
| *DNAH7* | rhesus | BS-F | GTTTGTTGTTTAGTAATAAAGGTTGT |
| *DNAH7* | rhesus | BS-R | TCACCTACTCACTACTCATAACTA |
| *DNAH7* | rhesus | SNP-F | CGGAACTGTGAGGATCTACTTCAG |
| *DNAH7* | rhesus | SNP-R | TGCTAGGTTTGGTTGTTATGACGA |
| *GLIS3* | chimp | BS-F | GTATATAGAGTAGTGGTTTGAGTTTT |
| *GLIS3* | chimp | BS-R | ACAACRAAACTAATCCTATAAAAAAC |
| *GLIS3* | chimp | SNP-F | CCCAGGTTTACGTGCTCCAGAGATG |
| *GLIS3* | chimp | SNP-R | CCCGACTTACTCGGGAGCGAGTT |
| *GLIS3* | rhesus | BS-F | GTAGAGGGGTATATAATTAGGGTAAG |
| *GLIS3* | rhesus | BS-R | TAAAACTTAAAAATACAACTACTACCCC |
| *HECW1* | chimp | BS-F | GGTTTTTTTTTTGTTTGTTTGTAAGT |
| *HECW1* | chimp | BS-R | AACAATTATATCCCATACAATACCC |
| *HECW1* | rhesus | BS-F | GYGTTTTAGGTTGATAATTGAATTT |
| *HECW1* | rhesus | BS-R | AACACTACCTACCRCCCCTT |
| *HECW1* | rhesus | SNP-F | TCCAGAGCTTTGGTTCACAAGG |
| *HECW1* | rhesus | SNP-R | CGGGAGGATAAAGCGATGAAGTG |
| *HTR5A* | chimp | BS-F | GGAGGTTGTGTTAGTTTTGGATT |
| *HTR5A* | chimp | BS-R | AAATAACAACRAAAAATACCCAAATAAAC |
| *HTR5A* | chimp | SNP-F | GAGGTTGCAAACATCCGGATT |
| *HTR5A* | chimp | SNP-R | CCACAGCTTCGGATATGGGT |
| *HTR5A* | rhesus | BS-F | GGAGTTTTGGGTTTTTGATAGTTTAGG |
| *HTR5A* | rhesus | BS-R | CCTATAAATTTAACCCCAATATCCTAC |
| *HTR5A* | rhesus | SNP-F | CCAAAACTGGCTTCCCTAGCAC |
| *HTR5A* | rhesus | SNP-R | CGATCATGACGTTGGAGATGCAC |
| *MCCC1* | chimp | BS-F | GYGATTATTTAGTATAGAGGTAGTT |
| *MCCC1* | chimp | BS-R | CTACTAAAACCRAACTCTAATCTTTA |
| *MCCC1* | chimp | SNP-F | GCCCGATGTAAACGGCTACGAG |
| *MCCC1* | chimp | SNP-R | GTATCCTGCTGTCCATCCGCATT |
| *MCCC1* | rhesus | BS-F | GTTTTTGATTGGTGTTTTTGGAGAT |
| *MCCC1* | rhesus | BS-R | CAACRACTCCCCAACACAAAAAC |
| *MCCC1* | rhesus | SNP-F | GACTCCCTCACATTCTTGGATCAG |
| *MCCC1* | rhesus | SNP-R | AGTCCTCAGAAATGGAAAACCGAA |
| *RHOBTB3* | chimp | BS-F | TTAGAGGGTTTTTTTTTAGGTAAGTG |
| *RHOBTB3* | chimp | BS-R | TAAAACAAAAACTAATACCCATCCTC |
| *RHOBTB3* | rhesus | BS-F | GGTAGTAGTAGTAGTAGGAGG |
| *RHOBTB3* | rhesus | BS-R | AAAAACAAACTACTACTACTCAAAAAC |
| *SCIN* | chimp | BS-F | TTGGAGGATTGAGAAGTTGGAGT |
| *SCIN* | chimp | BS-R | CCTACAAATCTCAAACCCCTAC |
| *SCIN* | rhesus | BS-F | AACACCAACTCCAACTTCTCA |
| *SCIN* | rhesus | BS-R | TTGGGTTTAGTTTAGTTATAAGGTTAG |
| *SCIN* | rhesus | SNP-F | GGGTAAGTCAATTGCGGACAAC |
| *SCIN* | rhesus | SNP-R | CTGGGTTTAGTCCAGCTACAAGG |
| *SORD* | chimp | BS-F | GTTTAGGTTGGTATAAAGGAGGA |
| *SORD* | chimp | BS-R | CCATAAAACTCTTTTAACTCTAAAATAC |
| *SORD* | rhesus | BS-F | TATGGAATTTTTTTGGTTTTTGGGT |
| *SORD* | rhesus | BS-R | CTATCACAAAAAAAAAAACCTAACCCC |
| *SORD* | rhesus | SNP-F | AACAAACAGCGTGCTACAGATGGA |
| *SORD* | rhesus | SNP-R | AGATCCCTGCCCTTTGCAACAG |
| *ST8SIA1* | chimp | BS-F | GTTTGTAAGTAAGAGAATTAAAGGTGTG |
| *ST8SIA1* | chimp | BS-R | TACCCTATTACAACACCCCCTAC |
| *ST8SIA1* | rhesus | BS-F | GGAGGGGGAGATTTTGATTTTTA |
| *ST8SIA1* | rhesus | BS-R | CTATCCTACCCCAAACTCCTC |
| *ST8SIA1* | rhesus | SNP-F | AGTCACGATCTATGGCCATGGT |
| *ST8SIA1* | rhesus | SNP-R | TTATCTCCAGGGCACGAGTGAG |
| *ZC3H12C* | rhesus | BS-F | GGGAGGTATGAGGGAGTTT |
| *ZC3H12C* | rhesus | BS-R | AACCTACTTACATTCTCTTCTCC |
| *ZFP90* | chimp | BS-F | GTTAGGGATTTTTGYGAGGGT |
| *ZFP90* | chimp | BS-R | ATTACTCCCTAAAACCAAAATTTTTC |
| *ZFP90* | rhesus | BS-F | GTTAAATTTAGGAGGGGTGT |
| *ZFP90* | rhesus | BS-R | TTAAACAACCCCTCAACAAATC |
| *ZNF396* | chimp | BS-F | GGGTTTATTTGGTAGGGTAGG |
| *ZNF396* | chimp | BS-R | CTCTCTAAAACCCCCAAACTAC |
| *ZNF396* | rhesus | BS-F | TTAAGAGGTTATAGGTAGTAAGGGT |
| *ZNF396* | rhesus | BS-R | CCTCTTATATCCCTTAAAATACCTAAC |

**Supplementary Table 5. Transmission of mutant *Impact* and *Slc38a4* alleles**

| ***Slc38a4*** cross | | Number of litters | WT offspring | Mut offspring | chi-square test |
|---|---|---|---|---|---|
| female | male | | | | |
| +/MT2AKO | +/+ | 4 | 14 | 17 | 0.590 |
| +/+ | +/MT2AKO | 8 | 19 | 15 | 0.493 |

| ***Impact*** cross | | Number of litters | WT offspring | Mut offspring | chi-square test |
|---|---|---|---|---|---|
| female | male | | | | |
| +/MTCKO | +/+ | 10 | 38 | 40 | 0.821 |
| +/+ | +/MTCKO | 9 | 29 | 28 | 0.895 |
